# MITA couples with PI3K to regulate actin reorganization during BCR activation

**DOI:** 10.1101/346445

**Authors:** Yukai Jing, Danqing Kang, Lu Yang, Panpan Jiang, Na Li, Jiali Cheng, Jingwen Li, Heather Miller, Boxu Ren, Quan Gong, Wei Yin, Zheng Liu, Pieta Mattila, Bing Yu, Chaohong Liu

## Abstract

As an adaptor protein, MITA has been extensively studied in innate immunity. However, its role in adaptive immunity as well as its underlying mechanism are not completely understood. We used MITA KO mice to study the effect of MITA deficiency on B cell development and differentiation, BCR signaling during BCR activation and humoral immune response. We found that MITA deficiency promotes the differentiation of marginal zone B cells, which is linked to the lupus-like autoimmune disease that develops in MITA KO mice. MITA is involved in BCR activation and negatively regulates the activation of CD19 and Btk and positively regulates the activation of SHIP. Interestingly, we found that the activation of WASP and accumulation of F-actin is enhanced in MITA KO B cells upon stimulation. Mechanistically, we found that MITA uses PI3K mediated by the CD19-Btk axis as a central hub to control the actin remodeling that, in turn, offers feedback to BCR signaling. Overall, our study has provided a new mechanism on how MITA regulates BCR signaling via feedback from actin reorganization, which may contribute to the effects of MITA on the humoral immune response.

## Introduction

MITA is expressed in hematopoietic cells in peripheral lymphoid tissues and is also highly expressed in non-lymphoid tissues such as the lung and heart. MITA locates to endoplasmic reticulum and mitochondria associated ER membrane [1]. MITA interacts with IRF3 to activate the type I interferon production in response to foreign DNA from a variety of intracellular pathogens [2, 3]. MITA has also been reported to be involved in the pathogenesis of systemic lupus erythematosus (SLE). The expression of MITA is low in SLE patients and MRL/lpr mice, of which the MRL/lpr mice have higher expression of JAK1 and lower expression of tyrosine phosphotases, SHP1 and SHP2 [4]. Furthermore, SLE B cells have highly activated BCR-, TLR7- and TLR9-mediated signaling pathways that are involved in reducing MITA expression [4]. The link between MITA and lupus demonstrates the importance of determining the regulation of MITA on BCR signaling after BCR activation.

PI3K is important for the development and differentiation of B cells and class I PI3K consists of a catalytic subunit-p110 and a regulatory subunit-p85 [5]. Splenic marginal zone (MZ) B cells and B1a cells are significantly reduced in p110δ deficient mice [6]. In B cells, the majority of cytosolic PI3K is associated with the membrane adaptor protein CD19, which effectively recruits p85α via tandem YXXM motifs [7]. Mice with B cell specific deletion of PTEN have elevated PI3K signaling as well as increased MZ B cells and B1a cells [8]. The absence of PTEN and the consequential increase in PI-3,4-P2 and PI-3,4,5-P3 can substitute the role of CD19 in promoting PI3K activity [8], and a similar function is also exhibited by another inositol phosphatase, SHIP. Mechanistically, CD19 phosphorylation activates Lyn that in turn recruits PI3Ks. Lyn activation also stimulates the activation of Vav and Tec, which contributes to PI3K activation in B-cells by activating Rac1, which then binds to PI3Ks via its RhoGAP domain [9, 10]. PI3K/PIP3 signaling then turns on Akt signaling via Akt /PDK-1 activation and reduces apoptosis by phosphorylating Foxo1/3 and promoting their nuclear export and degradation [11, 12], which thereby facilitates cell survival. PIP3 then binds and phosphorylates Akt, which activates mTORC1 directly or indirectly through the actions of the TSC1/TSC2 [13, 14]. MITA has been shown to regulate the tyrosine phosphatase, SHP [4], and whether it can regulate the inositol phosphatase SHIP during BCR activation still remains elusive.

BCR activation is vital for the function of B cells, such as the formation of germinal centers, isotype switching and somatic hypermutation. The initiation of BCR activation starts from the binding of antigen to the BCR, which is thought to induce the clustering of BCRs by cross-linking. The conformational change of the BCR exposes the activation site and recruits the recruitment and activation of Syk leads to the activation of Btk and PLCγ as well as further downstream signaling. Our previous research has shown that BCR activation leads to actin reorganization, which offers feedback to the BCR signaling by modulating the spatiotemporal organization of BCRs [15-19]. Btk activity leads to the activation of WASP, an actin nucleation factor, by regulating the activity of Vav, the generation of PtdIns-4,5-P2, and the direct phosphorylation of WASP [20]. Whether MITA regulates BCR signaling to orchestrate actin reorganization is unknown.

In our study, we used MITA KO mice to study the effect of MITA deficiency on BCR signaling and actin reorganization as well as to determine the underlying mechanisms. We found that the activation of the proximal positive BCR signaling molecule, CD19, and downstream molecule, Btk, was enhanced and that the proximal negative BCR signaling molecule, SHIP, was decreased in MITA KO B cells after BCR stimulation. The distal BCR signaling with PI3K-mediated Akt and mTORC1 activation was also upregulated as well as the phosphorylation of WASP and resultant actin reorganization. By using total internal reflection fluorescence microscopy (TIRFm), we found that the BCR clustering was reduced, but B cell spreading was increased in MITA KO B cells after stimulation with membrane associated antigens. Interestingly, the inhibition of PI3K rescued the defect of BCR clustering, enlarged B cell spreading as well as actin reorganization and the BCR signaling from the feedback of actin reorganization. Overall, our study provides a new regulatory pathway of BCR signaling based on the negative regulation of MITA on the PI3K central hub and regulation of actin reorganization via WASP.

## Results

### The deficiency of MITA alters the homeostasis of peripheral B cells, but not the developmental subsets in the bone marrow

In order to determine whether MITA affects the development of bone marrow B cells or not, we stained the different subpopulations of bone marrow B cells with BP-1 and CD24 antibodies to distinguish pre-pro, pro, early-pre, and B220-IgM antibodies to separate late-pre, immature and recirculating B cells. We did not observe any changes for most of the subpopulations except for decreased percentages and number of recirculating B cells in MITA KO mice (Fig 1A-C). We further examined the IL7R (CD127) expression that is crucial for the early development of bone marrow B cells, and not surprisingly we did not observe altered levels of CD127 in the MITA-deficient mice (Fig 1D). Therefore, MITA is dispensable for the development of B cells in the bone marrow. We further examined the deficiency of MITA on the differentiation of peripheral B cells. We used IgM-IgD antibodies to stain the T1, T2 and follicular (FO) B cells, CD21-CD23 antibodies to stain the marginal zone (MZ) B cells and CD95-GL7 antibodies to stain the germinal center (GC) B cells. Interestingly, we found that the percentage and number of MZ and GC B cells were significantly increased in MITA KO mice, but that of FO, T1 and T2 showed no changes (Fig 1E-L). We also did not find any difference for the proliferation and apoptosis of each peripheral subpopulation (data not shown). These results indicate that MITA suppresses the differentiation of MZ and GC B cells.

**Fig 1.**
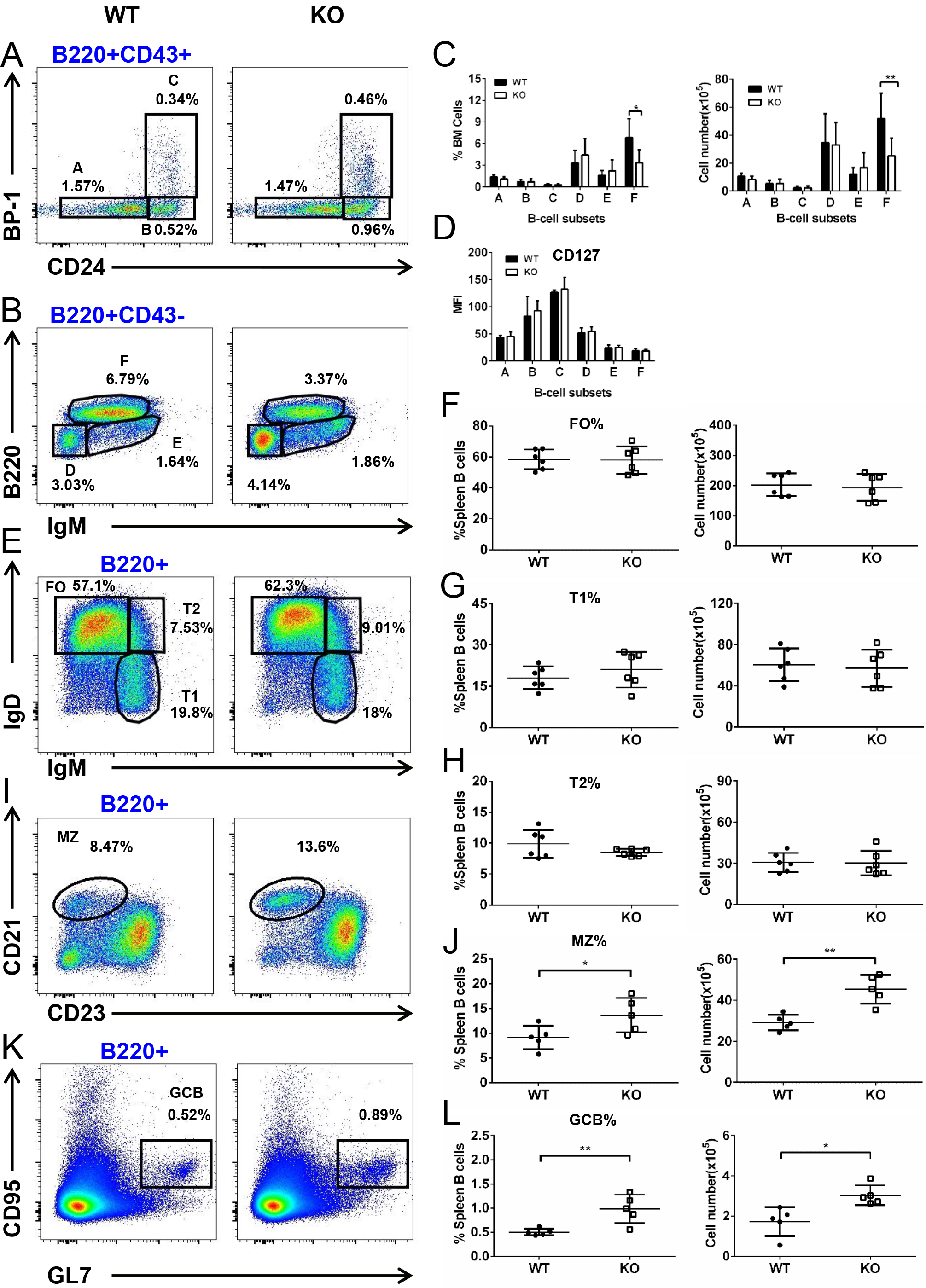
MITA deficiency has impact on the differentiation of MZ and GC B cells but not on the development of bone marrow B cells. Cells from BM of 6–8-week-old WT and MITA KO mice (n=6) were labeled with Abs specific for surface markers of pre-pro (A), pro (B), early-pre (C), late-pre (D), immature (E) and recirculating mature B cells (F), and analyzed by flow cytometry. Shown are representative dot plots (A and B), the average percentages (+SD) and numbers of total cells extracted from BM (C), and the average mean fluorescence intensity (MFI) of CD127 in B-cell subsets (D). Splenic cells (n=6) were stained with Abs specific for surface markers of transitional 1 (T1), transitional 2 (T2), follicular (FO), marginal zone (MZ) and germinal center (GC) B cells. Samples were analyzed using flow cytometry. Shown are representative dot plots (E, I and K), the average percentages (+SD) and numbers of subpopulations in spleen (F, G, H, J and L). *P < 0.05, **P <0.01.

### MITA is involved in BCR activation

Since we found that MITA was involved in the homeostasis of peripheral B cells, which is largely determined by BCR signaling, we next examined the involvement of MITA in BCR activation by confocal microscopy. WT splenic B cells were stimulated with soluble antigens (sAg) for varying lengths of time and then stained with antibodies specific for MITA. The spatiotemporal relationship between BCR and MITA was determined by the correlation coefficient to indicate the colocalization between BCR and MITA. We found that the correlation coefficient increased gradually up to 10 min and dropped afterwards (Fig 2A and 2B). Because the MITA KO is a germline deletion, we examined MITA expression in splenic B cells of MITA KO mice. We found that the signal of MITA expression was hard to be detected in KO B cells (Fig 2C). Since SLE can cause an abnormal increase of MZ B cells and MITA is linked to SLE [21-23], we examined the anti-double stranded DNA (dsDNA) antibody levels in WT and KO mice by ELISA. Interestingly, we indeed found an increase in anti-dsDNA antibody levels in KO mice (Fig 2E). Furthermore, we examined the architecture of the spleen of WT and MITA KO mice and correlating with the increased GC B cells in MITA KO mice, we found an enlarged and darker staining of the germinal centers (Fig 2D). These results imply that MITA is involved in B cell activation and controlling the peripheral tolerance of B cells.

**Fig 2.**
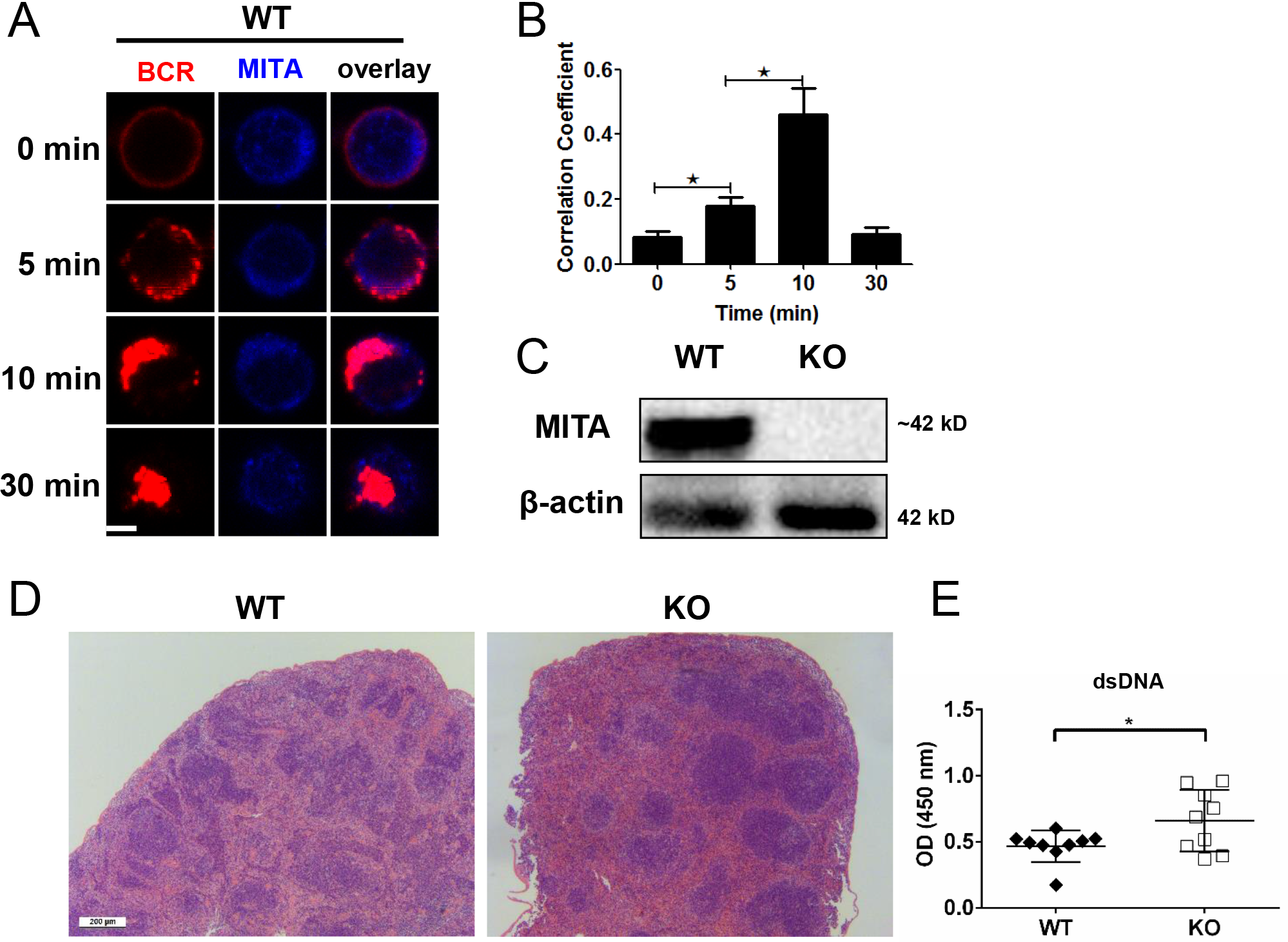
MITA is involved in the BCR activation. Purified splenic B cells from WT mice were labeled with AF546–mB-Fab–anti-Ig and activated by incubating with either streptavidin or the medium alone (0 minutes) as a control at 37°C for varying lengths of time. After fixation and permeabilization, samples were incubated for MITA and analyzed using confocal microscopy (CFM). Shown are representative images (A) and the correlation coefficients between the labeled BCR and MITA quantified using the ZEN 2.3 (blue edition) software (B). Scale bars, 2.5 μm. Splenic B cells from WT and MITA KO mice were lysed and probed with antibodies specific for MITA (C). Hematoxylin and eosin staining of spleen from WT and MITA KO mice (D). Scale bars, 200μm. ELISA quantification of anti-dsDNA Ab in the serum of WT and MITA KO mice (n=9). Dots represent individual mice (E). *P < 0.05.

### MITA deficiency upregulates the proximal positive BCR signaling

Since MITA is involved in the hyperactivation of B cells, we further investigated the effects of MITA on BCR signaling. Btk mediated PI3K signaling is critical for the maintenance and differentiation of MZ B cells, so we investigated whether this signaling axis could be augmented in MITA deficient B cells. First, we examined the effect of MITA deficiency on the spatiotemporal organization of phosphotyrosine proteins (pY), which represents the total levels of BCR signaling, and phosphorylated Btk by specific antibody staining using confocal microscopy. We found that the colocalization between pY/pBtk and BCRs were significantly increased in MITA KO B cells compared to that of WT B cells (Fig 3A-C). Furthermore, splenic B cells stimulated with sAg for varying lengths of time were lysed and probed with antibodies specific for pY and pBtk, then we found that levels of pY and pBtk were significantly enhanced in MITA KO B cells (Fig 3D-J). CD19 is an upstream regulator of Btk during BCR activation and similarly we examined the spatiotemporal relationship between BCRs and pCD19 using confocal microscopy and activation levels of CD19 using immunoblotting. Interestingly, we found upregulation of pCD19 by immunoblot as well as increased colocalization between BCR and pCD19 in MITA KO B cells (Fig 3E-H). All these results collectively suggest that MITA negatively regulates BCR signaling by inhibiting the activation of CD19 and Btk.

**Fig 3.**
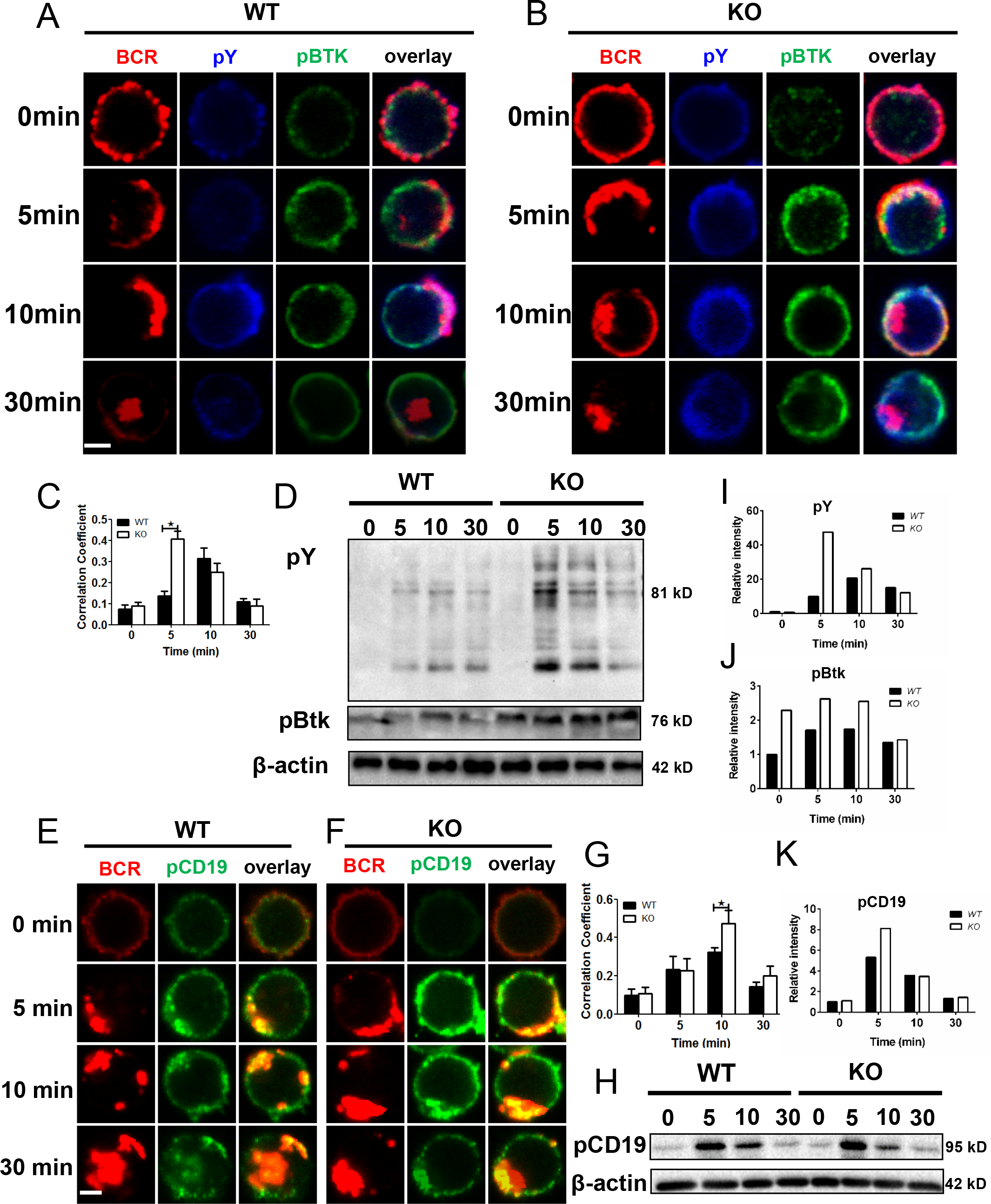
MITA deficiency up-regulates the positive BCR signaling-pY, pBtk and pCD19. Confocal analysis of splenic B cells from WT and KO mice labeled with AF546-mB-Fab–anti-Ig and then without or with streptavidin to activate. Cells were fixed, permeabilized, and stained with Abs specific for pY, pBtk (A and B) and pCD19 (E and F). Shown are representative images and the correlation coefficients between the labeled BCR and pY/pBtk (C) and pCD19 (G) from three independent experiments. Scale bars, 2.5 μm. Splenic B cells from WT and MITA KO mice were activated with Biotin-conjugated F(ab')2 Anti-Mouse Ig(M +G) plus streptavidin for indicated times. Cell lysates were analyzed using SDS-PAGE and Western blot, and probed for pY, pBtk (D) and pCD19 (H). β-actin as loading controls. Shown are representative blots of three independent experiments and blots’ relative intensity (I, J and K). *P < 0.05.

### MITA deficiency downregulates the activation of the negative BCR signaling molecule-SHIP

Since the activation of Btk and SHIP counters with each other, we examined the effect of MITA deficiency on the spatiotemporal organization between pSHIP and BCRs as well as the activation levels of SHIP. Splenic B cells were activated with sAg for varying lengths of time and then stained with pSHIP specific antibody for examination by confocal microscopy. We found that the correlation coefficient between BCR and pSHIP was significantly decreased in MITA KO B cells compared to that of WT B cells at 10 min (Fig 4A-C). Splenic B cells were also activated with sAg for varying lengths of time and then lysed and probed with antibody specific for pSHIP for analysis by immunoblotting. Interestingly, we found that the activation levels of SHIP were markedly decreased in MITA KO B cells at 5min after activation, the time of the peak intensity of pSHIP signal in WT cells (Fig 4D). These results indicate that MITA negatively regulates BCR signaling by enhancing the activation of SHIP.

**Fig 4.**
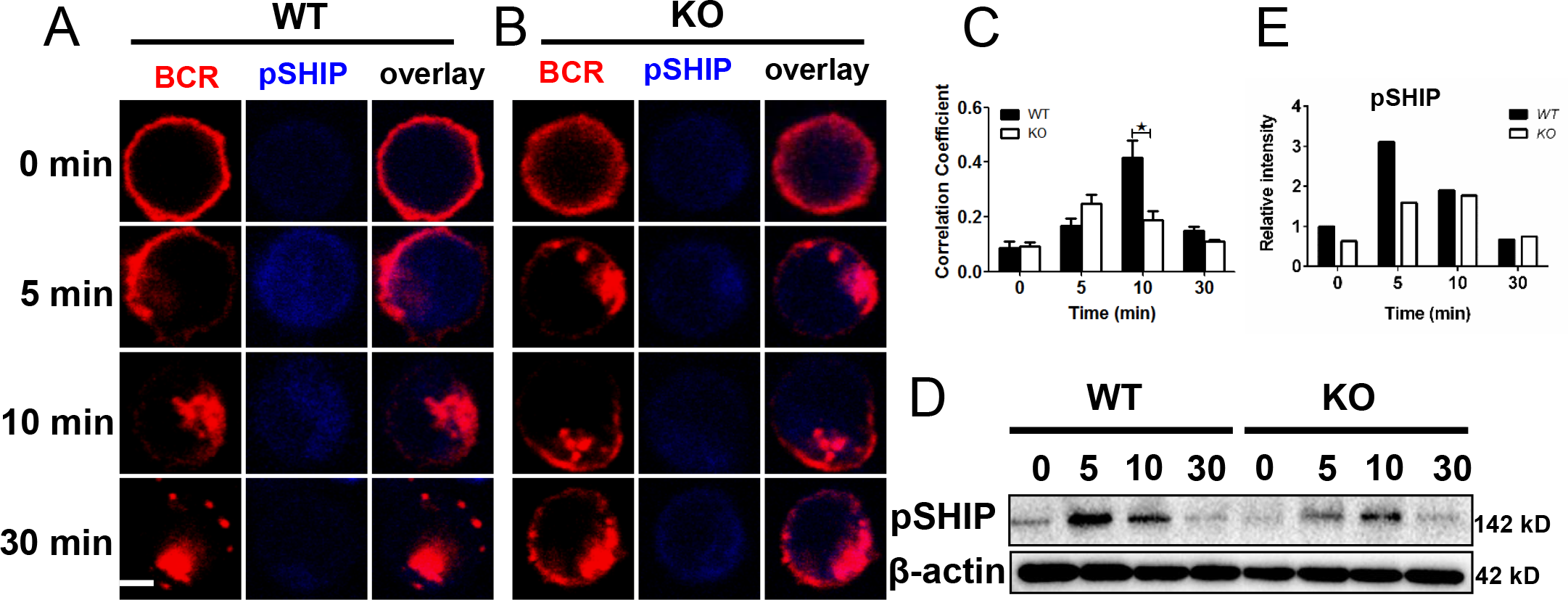
MITA deficiency down-regulates the activation of negative BCR signaling-SHIP. Confocal analysis of splenic B cells from WT and KO mice labeled with AF546-mB-Fab–anti-Ig and then without or with streptavidin to activate. Cells were fixed, permeabilized, and stained with Abs specific for pSHIP (A and B). Splenic B cells from WT and MITA KO mice were activated with Biotin-conjugated F(ab′)2 Anti-Mouse Ig(M +G) plus streptavidin for indicated times. Cell lysates were analyzed using SDS-PAGE and Western blot, and probed for pSHIP (D). β-actin as loading controls. Shown are representative images, blots, the correlation coefficients between the labeled BCR and pSHIP (C) as well as blots’ relative intensity (E) from three independent experiments. Scale bars, 2.5 μm. *P < 0.05.

### MITA deficiency causes an abnormal accumulation of F-actin via WASP

Our previous research has shown that Btk mediated PI3K signaling modulates the accumulation of actin via WASP [20]. So we investigated the effect of MITA deficiency on actin reorganization. Splenic B cells were stimulated with sAg for varying lengths of time and then stained with phalloidin and anti-pWASP antibody for evaluation using confocal microscopy. We found that the correlation coefficient between pWASP and BCRs was significantly increased in MITA KO B cells at 0 and 10 min (Fig 5A, 5B and 5E). Furthermore, we examined the upregulation of pWASP and accumulation of F-actin in a similar set up but using phosphoflow. We found that the pWASP levels were significantly higher at 5 and 10 min in MITA KO B cells upon stimulation compared to that of WT B cells (Fig 5C). The depolymerization phase of actin was attenuated in MITA KO B cells and caused an enhanced actin polymerization phase from 10 to 30 min (Fig 5D). These results indicate that MITA may negatively regulate the actin polymerization via inhibiting the activation of WASP, possibly via the Btk-PI3K signaling axis.

**Fig 5.**
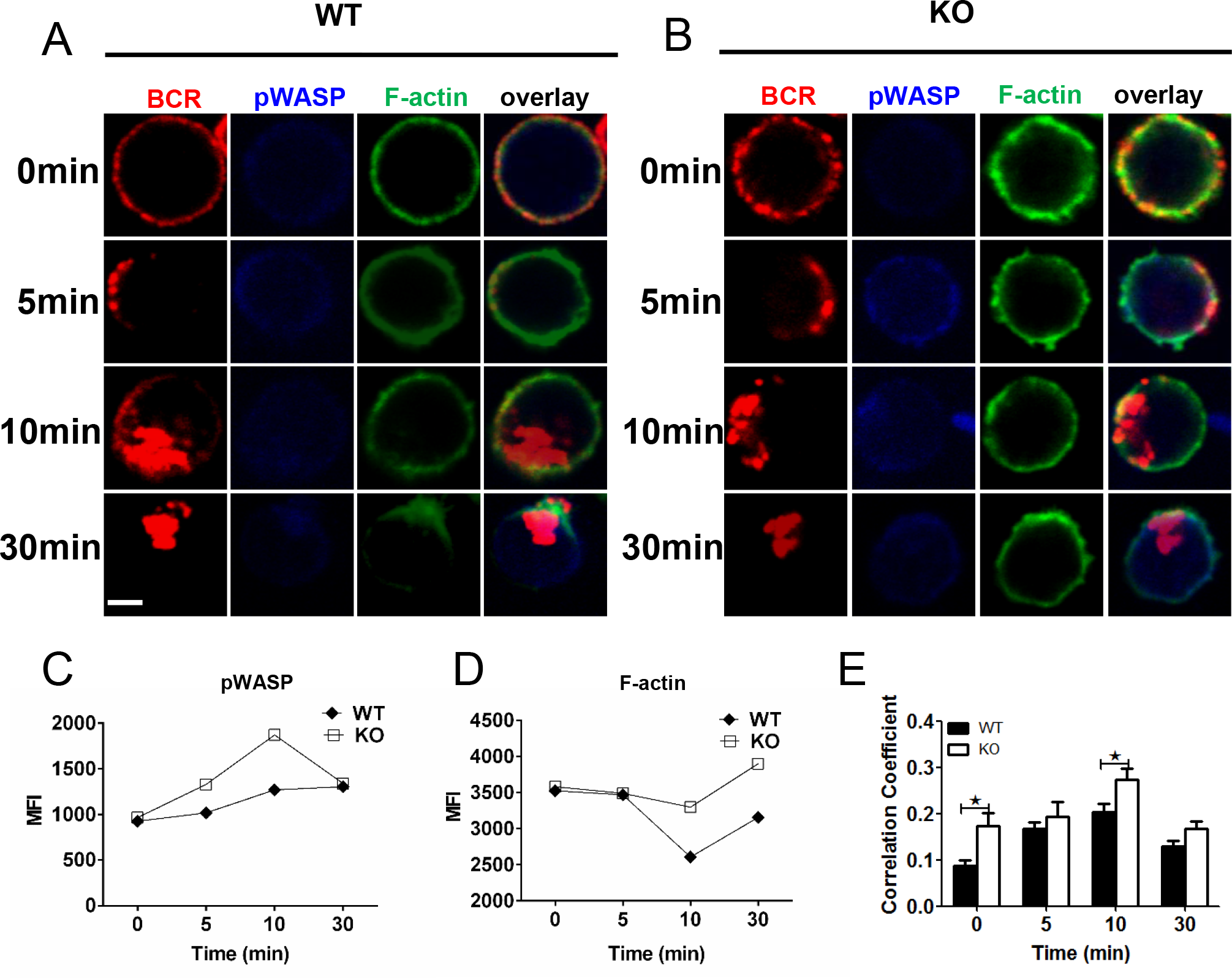
MITA deficiency up-regulates the actin polymerization via enhancing the activation of WASP. Confocal analysis of splenic B cells from WT and KO mice labeled with AF546-mB-Fab–anti-Ig and then without or with streptavidin to activate. Cells were fixed, permeabilized, and stained with Abs specific for pWASP and F-actin (A and B). Shown are representative images and the correlation coefficients between the labeled BCR and pWASP (E) from three independent experiments. Scale bars, 2.5 μm. *P < 0.05. Splenic B cells were labeled with anti-B220 and stimulated with sAg for indicated times and then stained with phalloidin and antibody specific for pWASP for phos flow cytometry. MFI of pWASP and F-actin in B cells was quantified using Flowjo software (C and D). Shown are levels of pWASP and F-actin from one of three independent experiments.

### MITA deficiency causes an abnormal accumulation of F-actin via PI3K mediated signaling

To further confirm whether MITA regulates actin reorganization via PI3K mediated signaling, we used a PI3K specific inhibitor, IC87114, to inhibit the activation of PI3K mediated signaling. In order to clearly observe the events occurring close to the membrane, we used total internal reflection microscopy to observe the accumulation of BCR accumulation, BCR signaling molecules and F-actin in the contact zone of B cells interacting with antigens tethered to lipid bilayers. We used membrane associated antigens (mAgs) to stimulate splenic B cells pretreated with or without IC87114 for varying lengths of time and then stained the samples with pY specific antibody and phalloidin. We found that BCR accumulation measured by MFI was significantly decreased in MITA KO B cells (Fig 6E), and that the BCRs could not form the central BCR cluster, which corresponds to downregulation of BCR signaling [17, 19], instead the KO displayed a punctate pattern (Fig 6A and 6C). However, the contact area of MITA KO B cells was significantly increased at very early time points indicating faster and more efficient activation (Fig 6F). The accumulation of pY and F-actin in the contact zone were also significantly enhanced in MITA KO B cells as compared to WT B cells (Fig 6G and 6H). WT B cells pretreated with PI3K inhibitor had significantly decreased BCR accumulation that was equal to the degree of that of KO B cells (Fig 6B and 6E). In KO B cells treated with PI3K inhibitor, the BCR accumulation was restored close to the degree of WT B cells (Fig 6D and 6E). For B cell spreading, WT B cells treated with PI3K inhibitor had B cell contact areas that were significantly decreased and equal to the degree of that of KO B cells (Fig 6F). In KO B cells treated with PI3K inhibitor, the contact area was almost restored to the level of WT B cells (Fig 6F). For pY signaling, WT B cells treated with PI3K inhibitor had BCR signaling that was significantly decreased compared to that of untreated WT B cells (Fig 6G). In KO B cells treated with PI3K inhibitor, the pY was partially restored (Fig 6G). For F-actin, WT B cells treated with PI3K inhibitor had their F-actin levels significantly decreased (Fig 6H). In KO B cells treated with PI3K inhibitor, the levels of F-actin were reduced close to that of WT B cells (Fig 6H). Furthermore, we investigated the effect of MITA deficiency on the PI3K signaling as well as signaling molecules downstream of PI3K, Akt and Foxo-1. The activation of Akt also leads to the initiation of mTORC1 signaling, so we also examined the activation of pS6 that is the immediate downstream signaling molecule of mTORC1. To this end, splenic B cells were activated with sAg, lysed and probed with antibodies specific for pPI3K, pAkt, pS6 and pFoxo-1. We found that the levels of pPI3K, pAkt, pFoxo-1 and pS6 were all clearly increased in MITA KO B cells (Fig 6I-N). All these results suggest that MITA uses PI3K as a central hub to regulate the actin reorganization which offers feedback to regulate the BCR clustering and BCR signaling.

**Fig 6.**
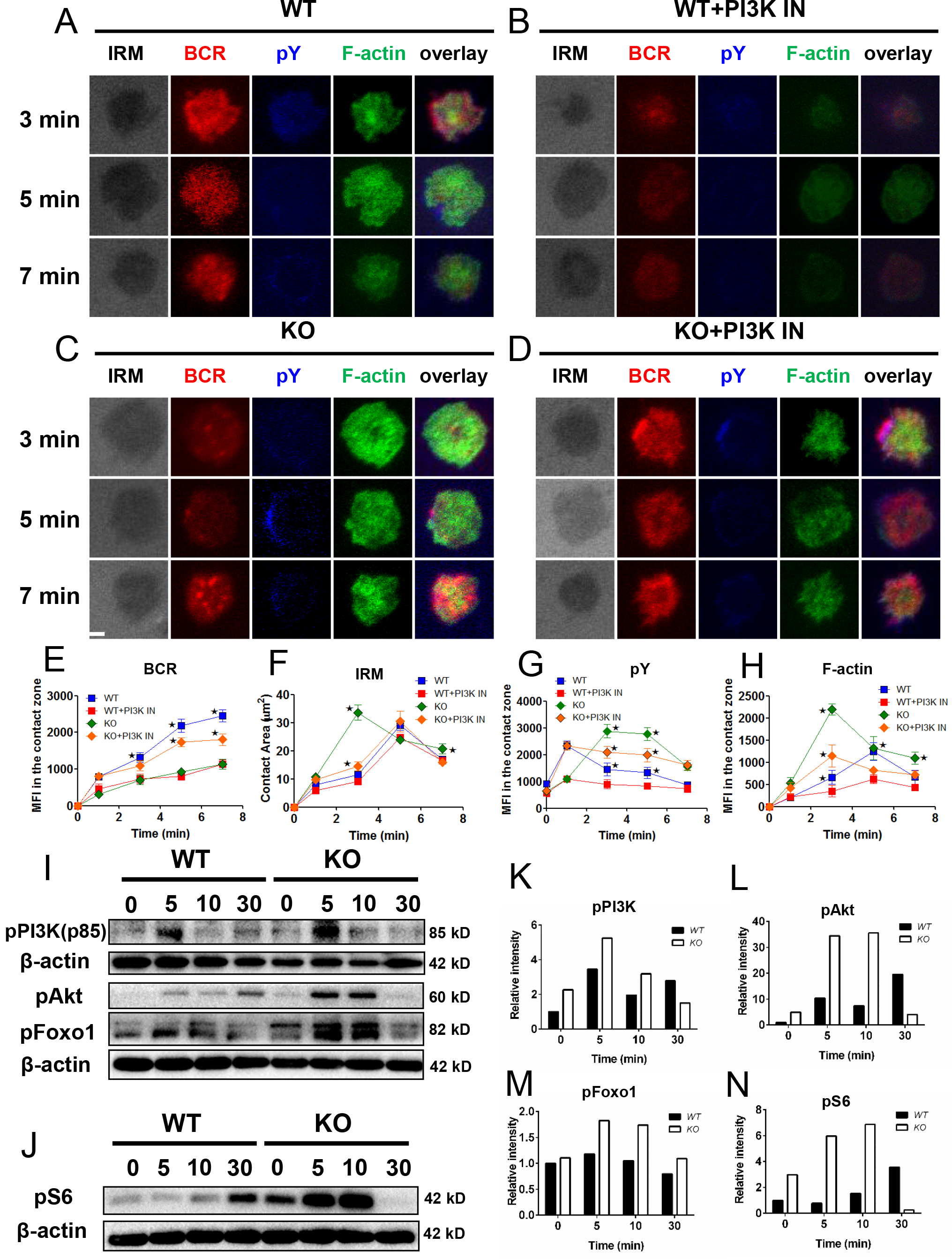
PI3K inhibition can rescue the abnormal accumulation of F-actin in MITA KO B cells. Splenic B cells from WT and MITA KO mice, pretreated with or without PI3K inhibitor, were incubated with AF546–mB-Fab–anti-Ig tethered to lipid bilayers with varying lengths of time, and then fixed, permeabilized, and stained antibodies specific for pY and F-actin. Cells were analyzed using TIRFm. Shown are representative images (A-D), the average values of the B-cell contact area (F), and the MFI of the pY (G), BCR (E) and F-actin (H) in the contact zone. Scale bars, 2.5 μm. *P < 0.05. Splenic B cells from WT and MITA KO mice were activated with Biotin-conjugated F(ab′)2 Anti-Mouse Ig(M +G) plus streptavidin for indicated times. Cell lysates were analyzed using SDS-PAGE and Western blot, and probed for pPI3K(p85), pAtk pS6 (I) and pFoxO-1 (J). β-actin as loading controls. Shown are representative blots from three independent experiments and blots’ relative intensity (K-N).

### MITA deficiency causes a reduced humoral immune response

To further investigate the effect of MITA deficiency on the humoral immune response, we immunized mice with NP-KLH to analyze the T-cell dependent immune response. After 14 days of immunization, the mice were euthanized and splenocytes and serum were harvested for further flow cytometry and serological analysis. After immunization, the frequency of FO B cells was decreased in both WT and KO mice but the absolute number of FO B cells was decreased in KO mice compared to that of WT mice (Fig 7A and 7F). Surprisingly, the number and percentage of MZ B cells were both decreased in KO mice compared with WT mice after immunization. Additionally, the number and percentage of MZ B cells were both decreased in KO mice after immunization compared to non-immunized (Fig 7B and 7G). The frequency and number of T1 cells were increased only in WT mice after immunization compared to non-immunized mice, and the frequency and number of T2 cells were increased both in WT and KO mice after immunization compared to non-immunized mice. The frequency of T2 cells were increased in KO mice after immunization compared to WT mice (Fig 7H and 7I). Surprisingly, the frequency and number of GC B cells were decreased in KO mice compared to WT mice after immunization, but the percentage and number of GC B cells were only increased in WT mice after immunization in contrast to non-immunized mice (Fig 7C and 7J). The percentage and number were only increased in PC B cells of KO mice and had no changes in MBC and PBC cells after immunization compared to WT mice (Fig 7D, 7E, 7K, 7L and Figure.EV1D). However, no difference for the frequency and number was observed for switched, unswitched and PBC between WT and KO mice after immunization (Fig.EV1A-D). Finally, we found that the titers of NP-specific IgG1 and IgM were both decreased in KO mice compared to that of WT mice after immunization (Fig 7N and 7O), which is consistent with the immunization results from DNA vaccine encoding ovalbumin [24]. All these results suggest that MITA is required for the efficient humoral immune response elicited by T cell dependent antigens.

**Fig 7.**
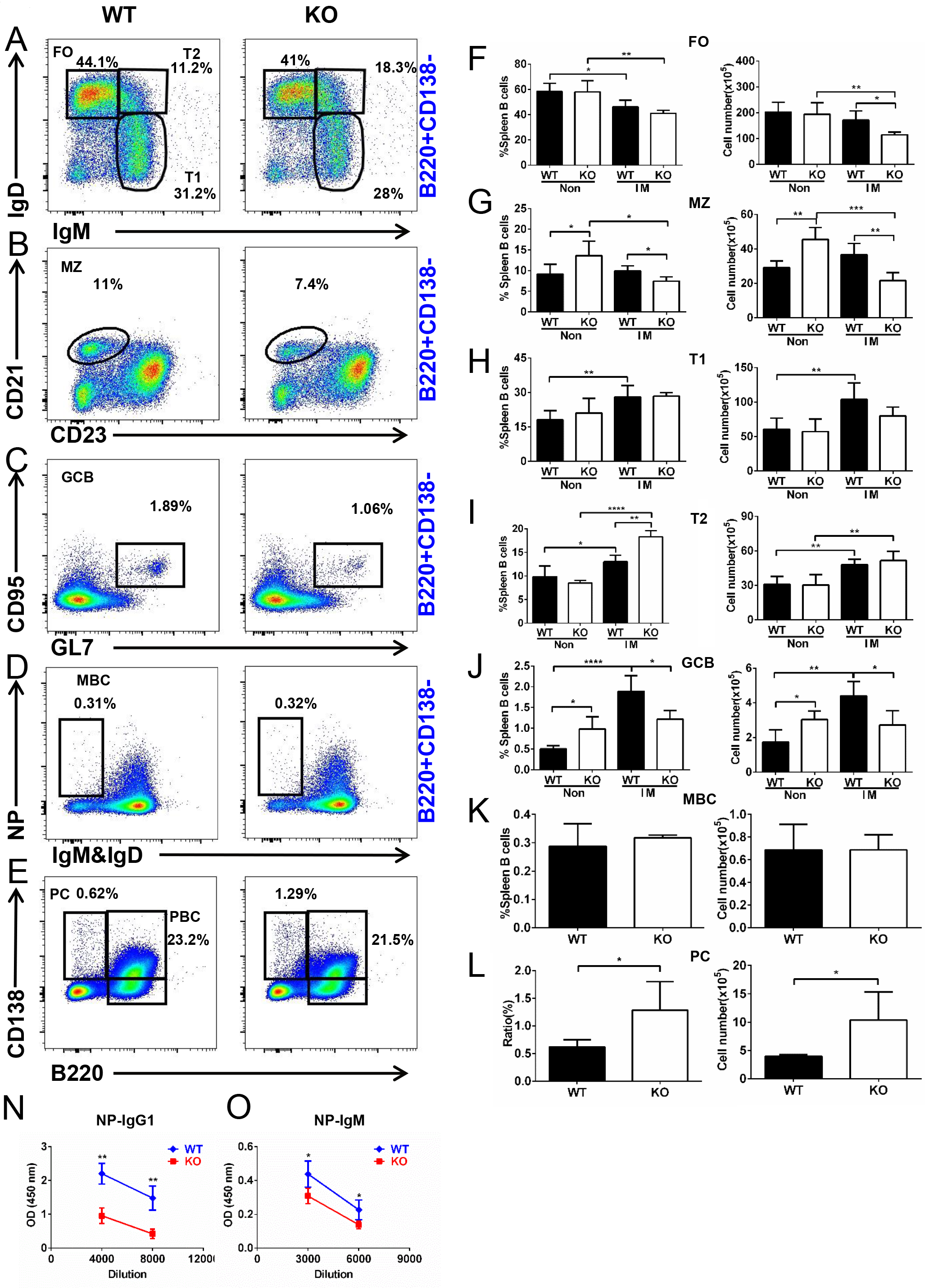
MITA deficiency reduces the T-independent Ab responses. Flow cytometry analysis of splenic cells from immunized WT and MITA KO mice (n=4) for follicular (FO) B, marginal zone (MZ) B, transitional 1 (T1) B, transitional 2 (T2) B, germinal center (GC) B, memory B (MBC) and plasma cell (PC) cells. Shown are representative dot plots (A-E), the average percentages (+SD) and numbers of subpopulations (F-L). ELISA analysis (n=4) of NP-specific IgG1 and NP-specific IgM in the serum (N and O). *P < 0.05, **P <0.01, *** p < 0.001 **** p < 0.0001.

### Discussion

In our study, we investigated the effect of the loss of MITA on BCR signaling and the underlying molecular mechanisms. We found that MITA negatively regulates BCR signaling by suppressing the CD19-PI3K-Btk proximal axis and the Akt-mTORC1 distal axis, and by enhancing the activation of SHIP, which also negatively regulates the actin reorganization. The inhibition of PI3K rescues the abnormal actin reorganization, which offers feedback to obtain the recovered BCR signaling and early B cell activation. Therefore, our study has provided a new regulatory mechanism of MITA on BCR signaling as well as actin reorganization.

We have shown that MITA can regulate the activation of WASP and resultant actin reorganization. It is the first time to link MITA to the actin cytoskeleton. Previous research demonstrated that MITA is correlated with the activation of autophagy that needs remodeling of the actin cytoskeleton [25]. MITA co-localizes with microtubule-associated protein 1 light chain 3 (LC3) and autophagy-related gene 9a (Atg9a) after sensing double stranded DNA [25]. But the underlying molecular mechanism is unknown. It would be interesting to examine whether MITA is involved in the BCR endocytosis that is critical to form the endosomal/lysosomal vesicles for autophagy. Our previous research has shown that the reduced BCR endocytosis upregulates the BCR signaling [26-28]. This may also contribute to the enhanced BCR signaling in MITA KO B cells. Another experiment to do is to inhibit the over-activated WASP and enhanced actin accumulation in MITA with *in vivo* treatments of Wiskostatin or Latrunculin B [15]. Additional experiments to perform may include possibly rescuing the MITA KO phenotype by crossing the MITA KO mice with WASP KO mice and then examining the early activation of these B cells, including BCR clustering and B cell spreading, as well as the differentiation of MZ B cells and lupus development. Based on what has been observed, it might be insightful to investigate whether MITA regulates other actin regulators, such as ezrin or Abp1.

Another remaining issue is how MITA regulates the expression level of CD19 as well as the phosphorylation of CD19. MITA mediates type I interferon immune response by acting as a direct DNA sensor and a signaling adaptor protein. MITA stimulates TBK1 activity to phosphorylate IRF3 or STAT6 upon stimulation with dsDNA. Phosphorylated IRF3s and STAT6s undergo dimerization, and then enter the nucleus to trigger the expression of genes such as IFNB, CCL2, CCL20 [29, 30]. It would be interesting to use a luciferase assay or CHIP-seq to identify whether the transcriptional factors, IRF3 and STAT6, can bind to the promoter of CD19 or not. Or we can induce bone marrow cells from MITA KO mice to express CD19 reporter with retrovirus in CD45.1 recipient mice to see if MITA regulates the expression of CD19 *in vivo* or not [31]. Another key point is that MITA mediated PI3K signaling is required to regulate the activation of WASP and the resultant actin reorganization. It would be interesting to treat the mice with PI3K inhibitor in vivo, or by crossing the MITA KO mice with PI3K mutant mice such as p110δ deficient mice, and examine the early activation of B cells and actin reorganization as well as the B cell differentiation and autoimmune phenotype in mice.

Another interesting finding of this study is the enhanced BCR signaling in MITA KO mice caused reduced GC formation and antigen specific antibody production. These results indicate that generation of GCs and antigen specific antibodies needs a specific window of BCR signaling. Too strong or too low of BCR signaling leads to the impaired generation of GCs and PCs, which causes lower production of antigen specific antibodies. All these results fit the findings from Dr.Westerberg’s group [32, 33]. Surprisingly, the percentage and number of PCs are increased in MITA KO mice, which is contradictory with the decrease of antibody production. It is possible that those PCs are not high affinity since the percentage and number of non-switched/switched PCs do not have changes. Sequencing somatic hyper mutation(SHM) is the best way to resolve this since it is not really possible to distinguish affinity from avidity by ELISA.

Overall, our study is the first to link the key adaptor protein-MITA, of the innate immunity, to its role in adaptive immunity. We found that it can modulate the BCR signaling via PI3K that regulates the actin reorganization via WASP. The regulation of MITA on BCR signaling leads to the control of humoral immune response. All of these findings will provide additional insightful thoughts for MITA studies in the innate immunity field as well.

## Materials and methods

### Mice

MITA KO mice on the C57BL/6 background were donated by Dr. Hongbing Shu’s lab in Wuhan University of China [34]. All mice were fed and maintained under the specific pathogen-free condition and mouse experiments were performed according to the guidelines of the Chinese Council on Animal Care and approved by the Ethics Committee of Animal Experimentation of Tongji Medical College (Wuhan, China).

### Purification of murine splenic B cells

Splenic lymphocytes were isolated by ficoll density centrifugation according to standard protocols. Splenic B cells were purified through deleting T cells by anti-Thy1.2 mAb (Biolegend) and guinea pig complement (Rockland Immunochemicals), and then incubated in Cell Culture Shelves for 1 hour.

### Preparation of monobiotinylated Fab’ Ab and Ag-tethered planar lipid bilayers

The planar lipid bilayer and liposomes were prepared as described previously^17^. The F(ab’)2 fragment (Jackson ImmunoResearch Laboratories) was used for making the monobiotinylated Fab’ fragment of anti-mouse IgM+G Ab (mB-Fab’-anti-Ig) following a published protocol [35]. To decrease the disulfide bond that links the two Fab’, 20 mM 2-mercaptoethylamine was needed, and the reduced cysteine was biotinylated by maleimide-activated biotin (Thermo Scientific). Next, Fab’ was purified, quantified and labeled with Alexa Fluor 546 (Thermo). To activate B cells with sAg, splenic B cells were labeled with AF546-mB-Fab’-anti-Ig (2μg/ml) and added in mB-Fab’-anti-Ig (8μg/ml) for 30 min and streptavidin (1μg/ml) for 10 min at 4°C. As a control, streptavidin was omitted. Cells were incubated at 37°C for indicated times. To activate B cells with mAg, cells were incubated with AF546-mB-Fab’-anti-Ig and mB-Fab’-anti-Ig tethered to planar lipid bilayers by streptavidin at 37°C for varying lengths of time. As a control, labeled B cells were incubated with transferrin (Tf)-tethered lipid bilayers where the molecular density of Tf was same as that of AF546-mB-Fab’-anti-Ig.

### Confocal and Total internal reflection fluorescence microscopy (TIRFm)

For confocal analysis, purified splenic B cells were stained with anti-CD16/CD32 mAb (BioLegend) to block Fcγ receptor (FcγR) and stimulated with AF546–mB-Fab’–anti-Ig with or without streptavidin (sAg) at 4°C. Cells were chased for 0, 5, 10, and 30 min at 37°C. After stimulation, cells were fixed with 4% paraformaldehyde (Thermo), permeabilized with 0.05% saponin (S4521-10G,sigma), and stained for anti-MITA (NBP2-24683SS, NOVUSBIO), anti-Phosphotyrosine (pY, 05-321, Merck Millipore), anti-pBtk (ab52192, abcam), anti-pCD19 (ab63443, abcam), anti-pSHIP1 (3941s, Cell Signaling Technology), AF488-phalloidin (R37110, thermofisher) and anti-pWASP (A300-205A, Bethyl) and analyzed using confocal microscope (Zeiss, LSM 780) with 405, 488, and 546 nm lasers. Colocalization was determined using the ZEN 2.3 (blue edition) software.

For TIRFm analysis, to image intracellular protein, B cells with or without PI3K inhibitor treatment were activated with an Ag-tethered lipid bilayer at 37°C for indicated times. Cells were then fixed, permeabilized, and stained for pY and AF488-phalloidin. Images were acquired using the TIRFm (Nikon Eclipse Ti-PFS). The B cell contact area was determined and analyzed using IRM images and NIS-Elements AR 3.2 software. MFI of each image in the B cell contact zone was determined using the same software. Background generated by Ag tethered to lipid bilayers in the absence of B cells controls was subtracted. For each set of data, more than 50 individual cells from three independent experiments were analyzed.

### Phenotype and intracellular protein analysis by Flow cytometry and phos flow

RBCs in collected bone marrow (BM) cells were lysed with ACK (Takara). BM cells and splenic cells were incubated with anti-mouse CD16/CD32. BM cells were stained with antibodies including APC-anti-CD43, PB-anti-IgM, PE-anti-BP1, PerCP-anti-B220, PE/Cy7-anti-CD24 and FITC-anti-CD127 (BioLegend). Splenic lymphocytes were stained with such antibodies as FITC-anti-CD19, FITC-anti-CD95, FITC-anti-Annexin V, FITC-anti-B220, Percp-anti-B220, BV510-anti-B220, PE-anti-CD23, Percp-anti-IgD, APC-anti-CD21, AF647-anti-GL7, PB-anti-IgM (BioLegend). For intracellular staining (PE-Cy7-anti-ki67, eBioscience), cells were fixed and permeabilized with Fixation/Permeabilization Kit (00-5123, 00-5223, eBioscience), washed with Permeabilization Buffer (00-8333, eBioscience). To analyze splenic MBC, PBC and PC B cells, other anti-mouse Abs also were used including PE-NP (Biosearch Technologies), BV510-anti-CD138 (Biolegend) and PB-anti-IgD (Biolegend).

For phosflow, primary B cells were incubated with Biotin-conjugated F(ab′)2 Anti-Mouse Ig(M +G) (115-066-068, Jackson ImmunoResearch) plus streptavidin at 37°C for varying lengths of time. Cells were fixed with Phosflow Lyse/Fix buffer, permeabilized with Phosflow Perm buffer III (BD Biosciences) and labeled with AF488-phalloidin and anti-pWASP. Cells were analyzed by LSRII multicolor flow cytometer (BD Biosciences CA, USA) and data analysis was performed using the FlowJo software (Tree Star, USA) as previously described [15].

### Western blot analysis

Purified B cells were activated with Biotin-conjugated F(ab′)2 Anti-Mouse Ig(M +G) plus streptavidin for indicated times. The cell lysates were separated by SDS-PAGE, transferred onto a nitrocellulose membrane and probed with anti-pY (05-321, Merck Millipore), anti-pBtk (ab52192, abcam), anti-pCD19 (3571S, Cell Signaling Technology), anti-pSHIP1 (3941s, Cell Signaling Technology), anti-pPI3K p85/p55 (4228s, Cell Signaling Technology), anti-pAkt (4060L, Cell Signaling Technology), anti-pS6 (4856S, Cell Signaling Technology) and anti-pFoxO1 (9461S, Cell Signaling Technology). Additionally, WT and KO splenic cell lysates were incubated with anti-MITA. **β**-actin was used to control the amount of total protein loaded. Immunoreactive bands were captured with the ChemiDoc™XRS + imaging systems (Bio-Rad). The levels were quantified by densitometry using Image Lab™ software (Bio-Rad), normalized against β-actin and expressed as fold increases over 0 min of WT.

### Immunization and enzyme-linked immunosorbent assay (ELISA)

For NP-KLH immunization, a mixture of NP-KLH/adjuvant was prepared according to the specification (S6322, Sigma) with a 0.2mg/ml concentration of NP-KLH (N-5060-25, BIOSEARCH). WT and MITA KO mice aged 6-8 weeks were injected subcutaneously with 200μl of the mixture. 2 weeks later, the spleen was harvested and splenic lymphocytes were collected with Ficoll for flow cytometry analysis. To measure the antibody levels of NP-specific subclasses, the serum was analyzed by ELISA using NP-bovine serum albumin (Biosearch Technologies)–coated plates and IgM and IgG1 specific secondary Ab (Bethyl Laboratories).

### Anti-double stranded DNA analysis

The serum levels of anti-dsDNA antibody were quantified by ELISA using the protocol as descripted [32].

### Treatment with PI3K inhibitor in vitro

B cells were pretreated with 3 μM of the p110δ specific inhibitor IC87114 (Calbiochem, Cat. No. 528118) for 30 min before and during the stimulation.

### Statistical analysis

Student t-test or the Mann–Whitney U test was carried out with Prism GraphPad Prism 6 Software (San Diego, CA) to assess the statistical significance. The difference was considered significant when p<0.05. *p < 0.05, ** p < 0.01, *** p < 0.001 **** p < 0.0001.

## Acknowledgments

This work was supported by Natural Science Foundation of China (81722002, 81861138002 and 31500709) and a start-up funding from Huazhong University of Science and Technology.

## Author Contributions

Y. Jing drafted the initial manuscript. C. Liu. and B. Yu designed the study, reviewed and revised the manuscript. Y. Jing, D Kang and L, Yang performed the confocal and Tirf experiments. Y. Jing, P. Jiang, N. Li, J. Cheng and J. Li carried out the flow cytometry assay, ELISA, and Western blotting. Y. Jing analyzed the data and generated figures. H. Miller, B. Ren, Q. Gong, W. Yin, Z. Liu and P. Mattila assisted with the manuscript. All authors approved the final manuscript as submitted and agreed to be accountable for all aspects of the work.

---

**Fig EV1.**
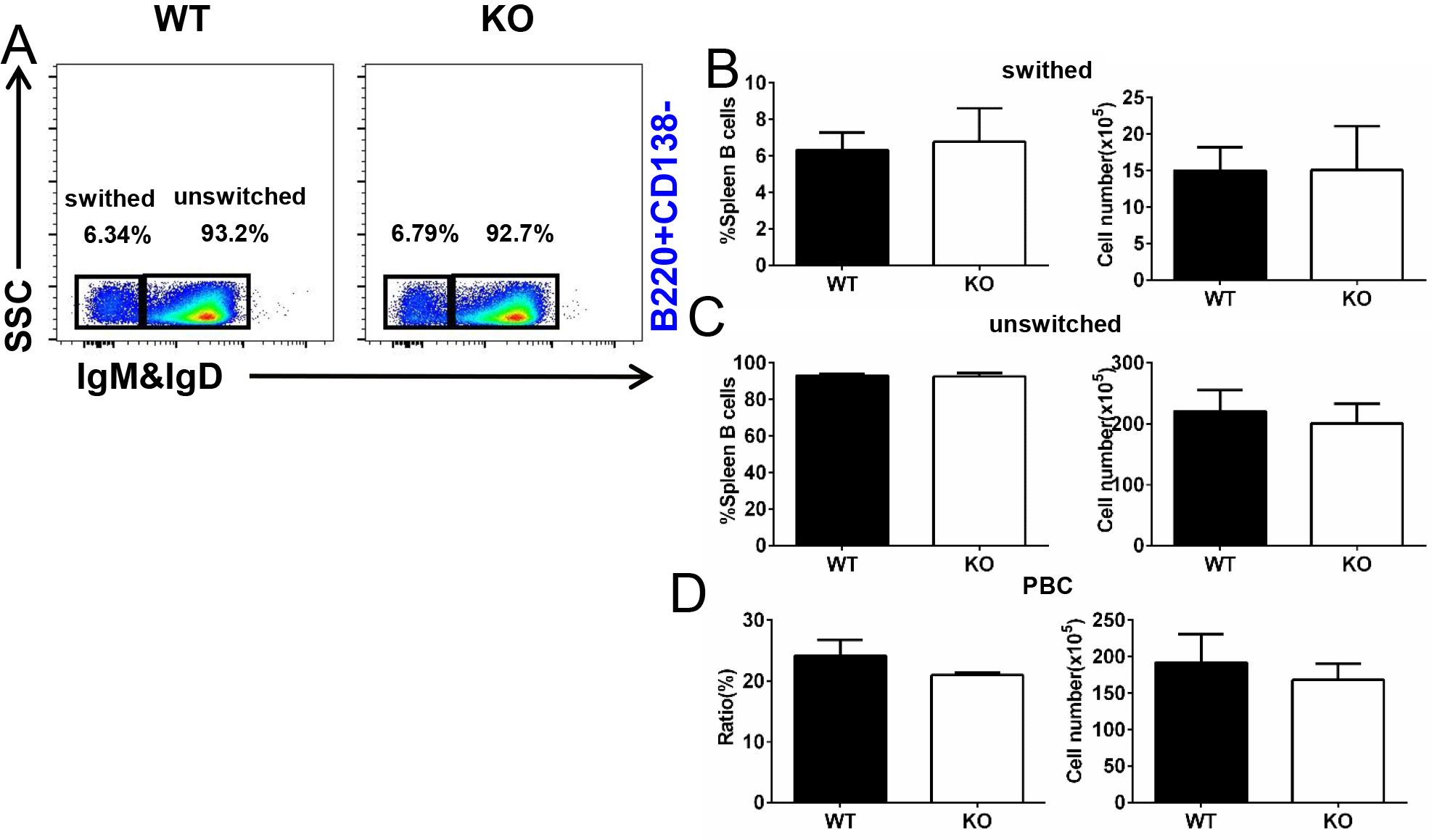
MITA deficiency has no impact on the differentiation of plasmablast B cells (PBC), switched and unswitched B cells. Flow cytometry analysis of splenic cells from immunized WT and MITA KO mice (n=4) for PBC, switched and unswitched B cells. Shown are representative dot plots (A), the average percentages (+SD) and numbers of subpopulations (B-D).

